# Scaling, correlations, and artifacts in biological timeseries

**DOI:** 10.1101/2023.07.10.548439

**Authors:** Denis F. Faerberg, Victor Gurarie, Ilya Ruvinsky

## Abstract

Time is a key variable in biology. Conceptual and methodological advances focused recent studies of timing in biological systems on monitoring trait dynamics in singled individuals. Here we contribute to this effort by analyzing general properties of individual timeseries. We make four broad claims. First, we show that for traits that range from behavior in animals, to growth in plants, to division timing in cell lineages, faster timeseries tend to be comprised of consistently shorter sub-stages, not a few unusually fast sub-stages. We demonstrate that this property constitutes a particular type of scaling that can be readily detected by a straightforward comparison of absolute and relative variability within timeseries data for any trait. We show that correlations within timeseries are necessary and sufficient for the type of scaling we describe and infer that the ubiquitous occurrence of scaling results from natural correlations within the continuous processes that govern trait dynamics. Second, failing to observe scaling indicates a break in correlation and, we argue, a switch in the mechanisms underlying the rate of trait change. Third, we caution that analyses of individual timeseries are highly susceptible to batch effects, but contend that it is possible to detect signal despite the artifactual correlations introduced by batching. Finally, we show that the timing of biological process that scale is overdispersed compared to matched but uncorrelated comparisons; this is an advantage in variable environments. Our results advance understanding of common properties of biological timeseries and offer practical tools for their further study.

## Introduction

Because processes along the entire continuum of life history – from growth and development to aging – unfold dynamically, considerations of time are ubiquitous in biology. However, the concept of “time” has several distinct meanings [1]. One is concerned with the relative order of events. This approach is exemplified by the heterochronic pathway, a cascade of genetic regulators that determine the order in which cell divisions and morphogenetic events occur during development [2]. A much-improved understanding of development now allows comparisons of the temporal order of events in embryogenesis of different species [3, 4], thus revisiting classical ideas, such as the “biogenic law” (for references see [5]), that promoted this view of biological time.

A different approach is to focus on measurements of absolute timing in biological systems. Both quantitative (e.g. size, number of cells, levels of gene expression, etc.) and qualitative (e.g. occurrence of a particular cell division, onset of adulthood, etc.) traits could be monitored over time to infer their dynamics (Figure 1A, left). Invariably, some individuals attain a certain value of a trait faster than others. An alternative to periodic monitoring of traits over time is to obtain the distribution of trait values in the population at a given time point (Figure 1A, right). Although less well-resolved than periodic monitoring, this snapshot approach nevertheless can quantify variation among individuals and determine whether average dynamics of a biological processes are similar between samples.

**Figure 1.**
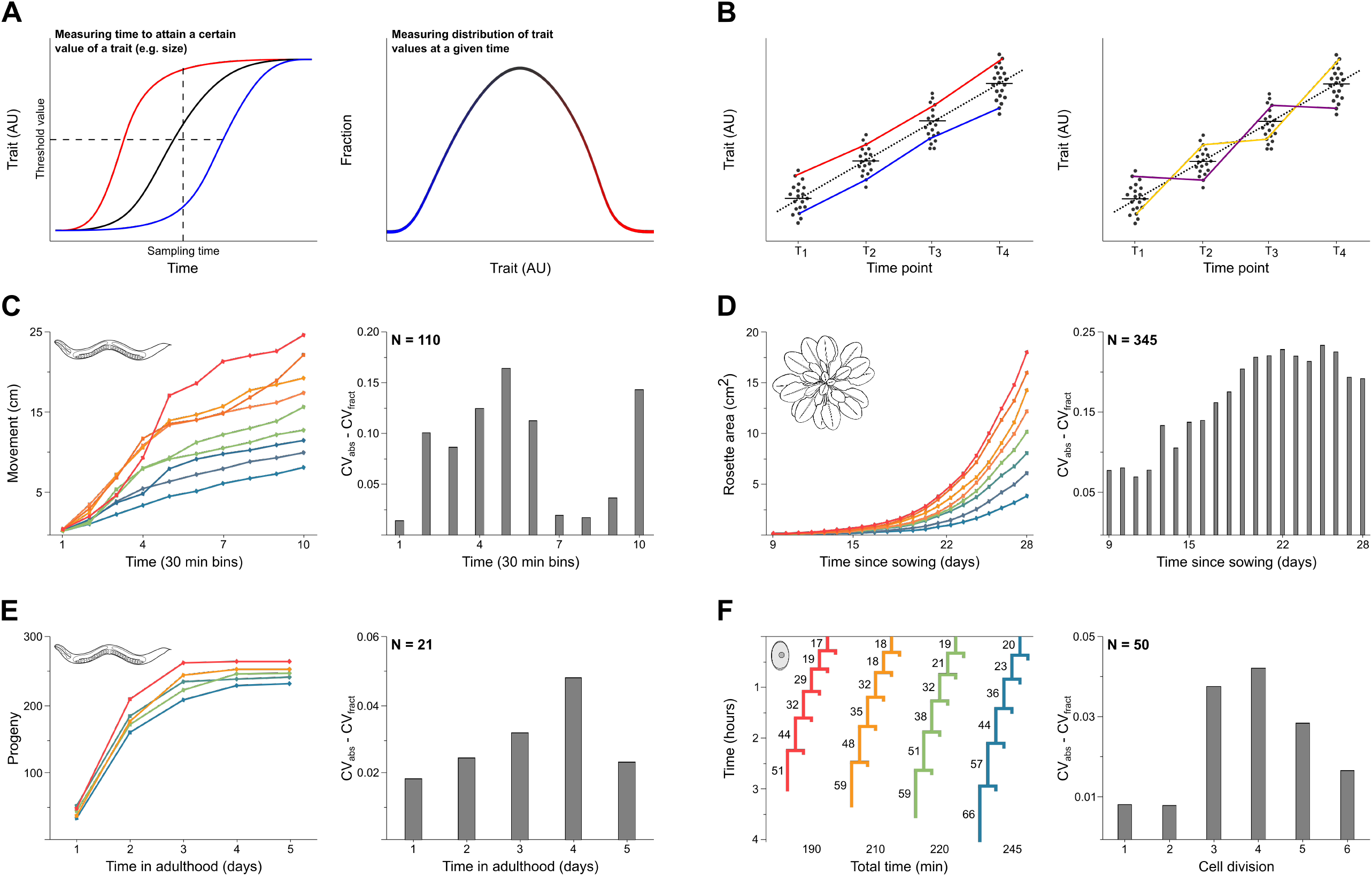
Absolute and fractional variability in biological timeseries. **(A)** Schematic representation of a quantitative trait changing over time (red – faster individuals, black – population average, blue – slower individuals). Dashed lines indicate two types of approaches to collecting data: measuring time when a certain value of a trait is achieved (horizontal) and measuring trait values at a certain time (vertical). The distribution of trait values at a given time is shown in the schematic on the right. **(B)** When traits are assessed in samples derived from populations at discrete times, it is not possible to distinguish whether some individuals are consistently ahead of others (red vs. blue in the left image) or whether distribution of trait values is due to inherent interindividual variability in rates of trait change (purple and orange in the right image). Panels **C-E** show change in traits in representative individuals at discrete times (left) and comparisons of absolute (CV_abs_) vs. fractional (CV_fract_) variability (right). **(C)** Distance traveled by individual *C. elegans* during a 5-hour period (measured over 10 consecutive 30 minute intervals) within the L3 larval period. **(D)** Daily increase in leaf area in *A. thaliana*. **(E)** Progeny produced by individual *C. elegans* mothers. **(F)** Timing of cell divisions (in minutes, indicated above branch bifurcations) leading to the ABalaaaal cell in four representative *C. elegans* embryos is on the left; absolute vs. fractional variability of division times of all 50 embryos is on the right.

The typical way in which experiments measuring dynamics of biological processes are carried out starts from establishing a synchronized population of individuals (one inherent challenge of this approach is discussed in the section “Batch effects inflate estimates of correlation within timeseries” below). Samples, drawn from such synchronized populations over time can be used to estimate mean trait values, variability within population, and approximate rate of trait change (Figure 1B). The latter inference necessarily relies on assumptions about the nature of the underlying processes. For example, sampling approaches make it impossible to distinguish between the following two scenarios. In the first, some individuals attain specific trait values consistently faster than others (Figure 1B, left). In the second, no individuals are consistently faster than others and the variability results from noise (Figure 1B, right). Distinguishing between these scenarios is essential for understanding the causes of interindividual variation. Therefore, although snapshot sampling can be convenient and reasonable in some experimental paradigms, it hampers quantitative descriptions of dynamic properties of biological processes.

The alternative approach to snapshot sampling from a common synchronized population is to follow individuals over time. Although this approach is not, strictly speaking, continuous (measurements are still made periodically), it allows inferences of interindividual variation and better inference of temporal dynamics of the measured processes. Over the last two decades, studies of individually-resolved timeseries provided insights into temporal variability of meiosis [6] and cell cycle [7] in yeast, mechanisms that regulate bacterial [8, 9] and animal [10] cell growth, govern dynamics of aging [11] [12] as well as growth and development [13-15] in multicellular organisms, and control behavioral individuality [16].

Individually-resolved biological timeseries contain more information than snapshot sampling from populations of individuals. Analysis of these data could offer novel insights into the regulatory mechanisms underlying the studied traits because it makes possible rigorous tests of precise hypotheses regarding relationships between different variables [17-19]. However, the use of such data also raises specific methodological issues. Here, we studied general statistical properties of individually-resolved timeseries.

## Results

### Widespread occurrence of scaling in biological timeseries

Several types of inferences could be made from individual-resolved timeseries, but not from snapshot population sampling. For example, larval development in the nematode *C. elegans* is divided into four stages, L1 through L4. Analysis of duration of larval stages inferred from individually-resolved timeseries revealed temporal scaling, that is, individuals with shorter overall larval development tended to have shorter L1, L2, etc. [14, 15, 20]. A more precise description of this kind of scaling is that interindividual variability of absolute duration (CV_abs_) of specific larval stages (e.g. L1) is greater than the variability of the fractions (CV_fract_) that these stages occupy in the entirety of duration of larval development (L1/ (L1+L2+L3+L4)). Because CV_abs_ and CV_fract_ can be straightforwardly calculated (template in Table S1) from timeseries data for any biological process, provided the data are individually-resolved, the quantity (CV_abs_ – CV_fract_) may be a useful indicator of whether the underlying process scales.

While the example above involves duration of larval stages, using the same rationale, it is possible to study scaling of any quantitative trait (height, volume, number of cells, etc.) that changes over time. To ascertain the generality of our approach, we tested whether the scaling seen for timing of larval development in *C. elegans* was a unique property of this process in this species. We first considered a different trait in the same species. Having monitored exploratory movement of over 100 worms from L1 to adulthood, Stern *et al*. observed behavioral individuality, that is, some animals moved consistently more than others [16]. This phenomenon is illustrated in Figure 1C that shows cumulative movement by nine representative worms over 300 minutes during the L3 larval stage. As discussed in the paragraph above, scaling is inferred when CV_abs_ exceeds CV_fract_. In a sample of the worms studied by [16], over 10 nonoverlapping 30-minute intervals, CV_abs_ was greater than CV_fract_ (Figure 1C), revealing scaling.

We next considered a different kind of trait in a different species – vegetative growth, estimated from the leaf rosette area in the thale cress *A. thaliana*. Several hundred individual plants were measured daily for approximately three weeks [21]. As was the case for the durations of larval stages [14, 15, 20] and exploratory activity (Figure 1C) in *C. elegans*, plant growth scaled over time (Figure 1D; analysis of growth under alternative conditions that yielded the same inference of scaling is shown in Figure S1A). Another quantitative trait we examined – offspring production in *C. elegans* (Figure 1E) – also scaled over time.

Finally, we tested whether timing of discrete developmental events scaled over time. Embryonic cell lineages from multiple individuals have been determined with great temporal precision in *C. elegans* [22-24] and the ascidian *P. mammillata* [25]. Comparison of CV_abs_ and CV_fract_ of time intervals between sequential divisions in a given lineage scaled in both *C. elegans* (Figure 1F, another lineage analyzed in Figure S1B) and *P. mammillata* (Figure S1C, S1D). Taken together, comparisons of CV_abs_ and CV_fract_ across different traits in highly divergent species suggested that scaling is common in biological timeseries data.

### Correlations within timeseries are necessary and sufficient for scaling

To better understand whether the observation of pervasive scaling was expected, we sought to derive analytical relationships between quantities that could be readily measured in biological experiments to infer scaling. In particular, we wished to examine the relationship between CV_abs_ and CV_fract_ (for ease of notation: *c*_*i*_ and *r*_*i*_, respectively).

Assume timeseries data that consist of positive and random variables *t*_*i*_ that are the durations of “stages”, where *i* = 1, 2, …, *N*. Coefficients of variation of stage durations are defined by

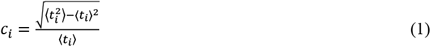

where ⟨ ⟩ denote averaging over many realizations of random variables *t*_*i*_. Fractional stage durations are defined as

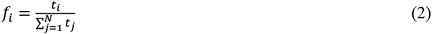

Coefficients of variation of fractional stage durations can be expressed as

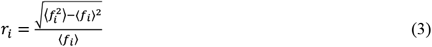

Previously, we demonstrated [20] that as long as fluctuations of *t*_*i*_ are much smaller than their average, 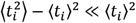, the following relationship holds true

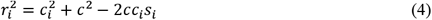

Here, *c* is the coefficient of variation of the total time 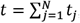, defined in the same way as its counterparts *c*_*i*_. In other words

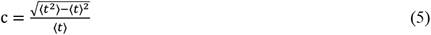

*s*_*i*_ are correlation coefficients between the duration of a stage and the total duration defined by

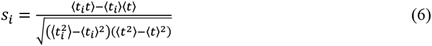

To understand the relationship between the coefficients of variation *r*_*i*_ and *c*_*i*_ we introduce the basic correlation coefficients *K*_*ij*_ between the durations of the stages

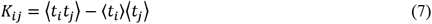

We denote the average of each stage by *a*_*i*_ = ⟨*t*_*i*_⟩, while the average of the total duration of all stages is 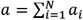. The correlation coefficients *K*_*ij*_ are the most basic quantities describing correlations among stage durations *t*_*i*_. All other quantities introduced earlier can be expressed in terms of these correlation coefficients. In particular,

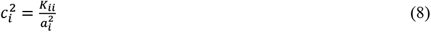

and

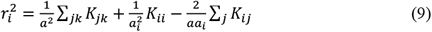

Consequently,

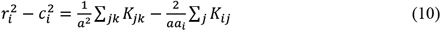

Multiplying by *a*_*i*_ and summing allows us to make the following observation

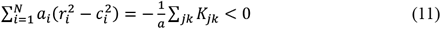

While the equality above is straightforward, the inequality follows from the fact that ∑_*jk*_ *K*_*jk*_ is the square of the standard deviation of the total duration *t*,

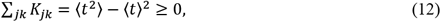

which is nonnegative by definition and can only be zero if the total duration *t* is not random. The inequality 11 is instructive since it indicates that *c*_*i*_ tends to be greater than *r*_*i*_, that is, CV_abs_ – CV_fract_ > 0.

Although the relationship between CV_abs_ and CV_fract_ derived above was consistent with empirical observations (Figure 1), we wished to better understand its implications. In particular, under what circumstances is *r*_*i*_ (CV_fract_) minimized, which could increase CV_abs_ – CV_fract_, indicating greater scaling? If durations of stages (*i* = 1, 2, …, *N*) are proportional to each other, *t*_*i*_ = λτ_*i*_ (where τ_*i*_ are fixed nonrandom time intervals and λ are random coefficients identical for all stages), then fractional durations *f*_*i*_ (Equation 2) remain constant from one individual to another and *r*_*i*_ = 0. This situation would constitute perfect scaling. We concluded that at least some scaling is expected in biological timeseries (Equation 12) and the greater the correlation between stage durations, the greater the extent of scaling. The derivations above were obtained for durations of stages in a continuous process (larval or other life history periods, time between consecutive cell divisions, etc.), but the same calculations also apply to measurements of physical quantities (e.g. size) that change over time.

### Lack of scaling is an indicator of change in the underlying processes

As shown above, CV_abs_ (*c*_*i*_) is typically greater than CV_fract_ (*r*_*i*_) in biological timeseries, as is expected from the mathematical relationship of these two quantities. To investigate the conditions when this relationship is reversed, we focused on the quantity 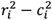 (see Equation 10) that is greater than zero when *r*_*i*_ > *c*_*i*_. The relationship between *r*_*i*_ and *c*_*i*_ was derived assuming that *r*_*i*_ ≪ 1 [20]. Therefore, an analytically rigorous attempt to maximize 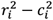 would go beyond the formalism used here. This may not be a practical problem, however, because in the biological situations we have analyzed, *r*_*i*_ ≪ 1. As a reasonable approximation, we investigated the heuristic criteria for *r*_*i*_ to become greater than *c*_*i*_. Equation 10 can be rewritten in the following way

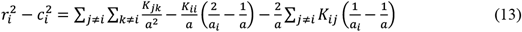

The first of the three terms on the right side of Equation 13 is nonnegative (compare with Equation 12). The second term is nonpositive, since 1/*a*_*i*_ > 1/*a* and *K*_*ii*_ ≥ 0. The third term could be of either sign. To make 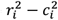 positive (i.e., *r*_*i*_ > *c*_*i*_), which is to say CV_abs_ < CV_fract_, *K*_*ii*_ should be as small as possible, while *K*_*ij*_ should equal zero or be negative. This would mean that the duration of the *i*-th stage is either not correlated or negatively correlated with the durations of other stages, but the durations of all other stages are positively correlated to each other.

Examples of biological processes that satisfy these conditions on correlations between stages do exist. Uppaluri *et al*. have monitored growth of individual *C. elegans* larvae for approximately 60 hours from hatching to adulthood [26]. At the end of this observation period, inter-individual variability in volume was over two-fold [26]. We divided individual 60-hour recordings of growth into six equal 10-hour periods and found that growth scaled for all except for the first period, the one roughly corresponding to the L1 stage (Figure 2A). Using different data and different measurement strategies, two other groups independently concluded that growth during the L1 does not follow the same patterns as growth during the other three larval stages [14, 27].

**Figure 2.**
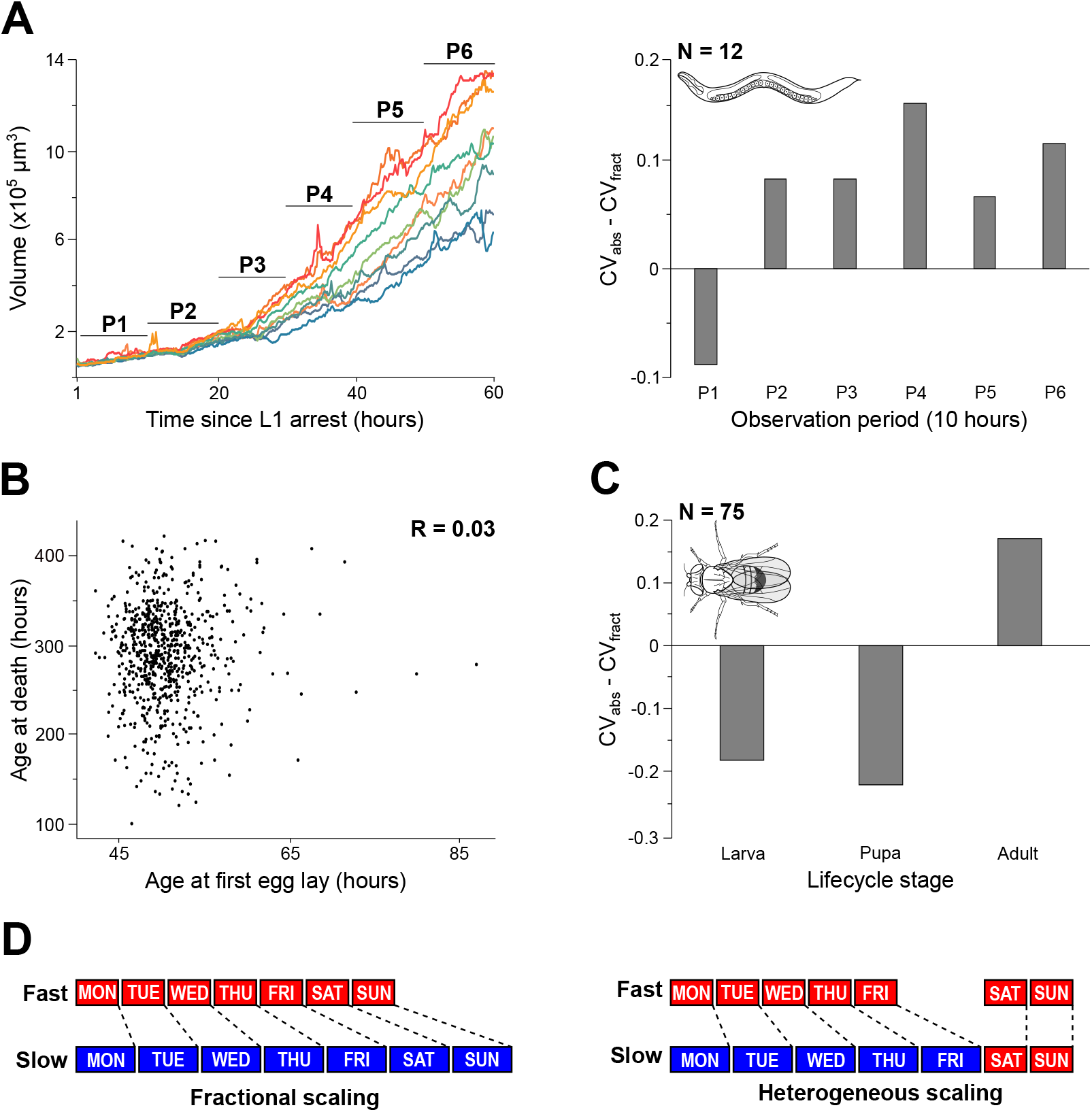
Increased fractional variability results from low correlations. **(A)** Growth curves of nine representative *C. elegans* larvae (left). The entirety of the observation period was divided into six 10-hour intervals. Absolute variability (CV_abs_) of volume gain is greater than fractional variability (CV_fract_) in all periods, except in P1 (right). **(B)** Correlation between the age at the onset of reproduction and lifespan in *C. elegans* (N=723). **(C)** In *D. melanogaster* males, absolute variability (CV_abs_) is greater than fractional variability (CV_fract_) for the duration of adulthood, but not for the larval or pupal periods. **(D)** Illustration of fractional vs. heterogeneous scaling. Sizes of analyzed samples are shown in the top left corners of the corresponding bar plots (**A, C**).

Analysis of Equation 13 reveals that the observation of CV_abs_ – CV_fract_ < 0 during a particular period within a biological process is an indication of the lack of correlation between the event(s) occurring during this and other periods. Despite correlations being common in biological timeseries (Figure 1), breaks in correlation also occur. For example, in *C. elegans*, the age at first reproduction is not correlated with that of the individual’s remaining lifespan (Figure 2B) [11]. Similarly, our analysis revealed the lack of uniform scaling between larval, pupal, and adult stages of *D. melanogaster* life history (males in Figure 2C, females in Figure S2A), fully consistent with the finding of little to no correlation between durations of these stages [13]. Duration of larval and pupal development are not correlated in two species of carrion blowflies, *L. serricata* and *C. vicina* [28] (Figure S2B, C).

The fact that durations of some periods of a biological timeseries are correlated, while others are not, may offer insights into the nature of the underlying processes. The observation of a correlation between duration of periods P_1_ and P_2_ could plausibly be explained by the same mechanism determining how long these periods last. If the length of period P_3_ is not correlated with P_1_ or P_2_, its duration may be determined by a different mechanism. The break in correlation does not have to be abrupt. For example, in bacteria, correlation in protein production between a cell and its descendants gradually decays over several generations [8] and there is some, albeit modest, correlation between the rates of growth of mother and daughter cells [9]. Exploratory movement in *C. elegans* individuals is highly correlated on the timescales of a few minutes, but decays considerably over times of approximately one hour (Figure S3).

The understanding that correlation between some, but not other processes could reflect changes in the underlying mechanisms could be consequential. For example, in contrast to the tight regulation of embryonic development that may impose limits on its temporal variation, aging appears to be dominated by stochastic processes [11, 29]. A realization that individual-specific responses to variable environment largely determine variability of lifespan [30], compels the search for biomarkers of individual longevity [31].

A schematic illustration of the change in scaling (i.e., correlation between some, but not other processes) is shown in Figure 2D. In the left scenario, all seven periods scale between “Fast” and “Slow” individuals according to the same underlying rule. In contrast, in the scenario on the right, whereas one rule governs proportional differences of the first five periods, the last two periods are not lengthened according to the same rule. In practice, while the nature of the underlying biological processes is almost never known, computing correlations between readily measurable quantitative variables (duration of developmental stages, number of cells, organ size, etc.) is relatively straightforward. For this reason, the observation of a breakdown in scaling, i.e., CV_abs_ < CV_fract_, could serve as an easily obtainable indicator of change in the underlying process and as a point of departure for in-depth functional studies.

### Batch effects inflate estimates of correlation within timeseries

The relevance of observing correlations (Figure 1) or their occasional lack (Figure 2) in biological timeseries is predicated on one critical assumption – that these statistical inferences reflect salient features of the underlying biological processes. However, certain features of experimental design can create an appearance of correlation in data series that, in reality, are correlated weakly or not at all. Whereas larger sample sizes are generally desirable, in practice they may be difficult to obtain due to limited sample availability, laborious experimental design, etc. In such cases, it is common to process samples in multiple batches and then either pool them together for a combined analysis or to use batch means to estimate trait values in the whole population. This approach can be powerful, particularly if mean values are considered. However, if there are systematic differences between batches that are due to causes extrinsic to the studied system, pooling batches together would artificially inflate correlations. This may be a relatively common phenomenon.

The inference of scaling in the timing of cell divisions during *C. elegans* embryonic development (Figure 1F) was made using 50 embryos. These data were obtained from five different strains (Table S1), 10 embryos per strain [23, 24]. Each set of 10 embryos could be considered as a batch – although the relative timing of cell division events was similar across the five strains, systematic differences were evident (Figure 3A, Figures S4A, B). Using a different approach, a recent study also detected batching in these data [32]. However, underscoring the quality of these data, straightforward linear transformations made the estimates of average cell division timing obtained from the five strains almost perfectly coincident (Figure 3B). The temporal precision of cell division events during *C. elegans* embryogenesis was faithfully captured by the highly accurate recordings despite batches differing by factors of ∼6-11% that were presumably due to strain-intrinsic or extrinsic environmental causes. To test whether the scaling inferred for the entire set (N=50; Figure 1F) could also be inferred from each of the smaller (N=10) batches, we repeated the analysis in Figure 1F separately for each of the five strains. The timing of the last four of the recorded divisions, scaled in all five strains (Figure 3C). For the first two divisions (ABa and its daughter), the evidence from the five strains was somewhat discordant. It is not currently clear whether the apparent difference between the first two and the subsequent four divisions was due to small sample sizes or to different mechanisms that regulate the timing of cell division during these two periods, as argued in the previous section.

**Figure 3.**
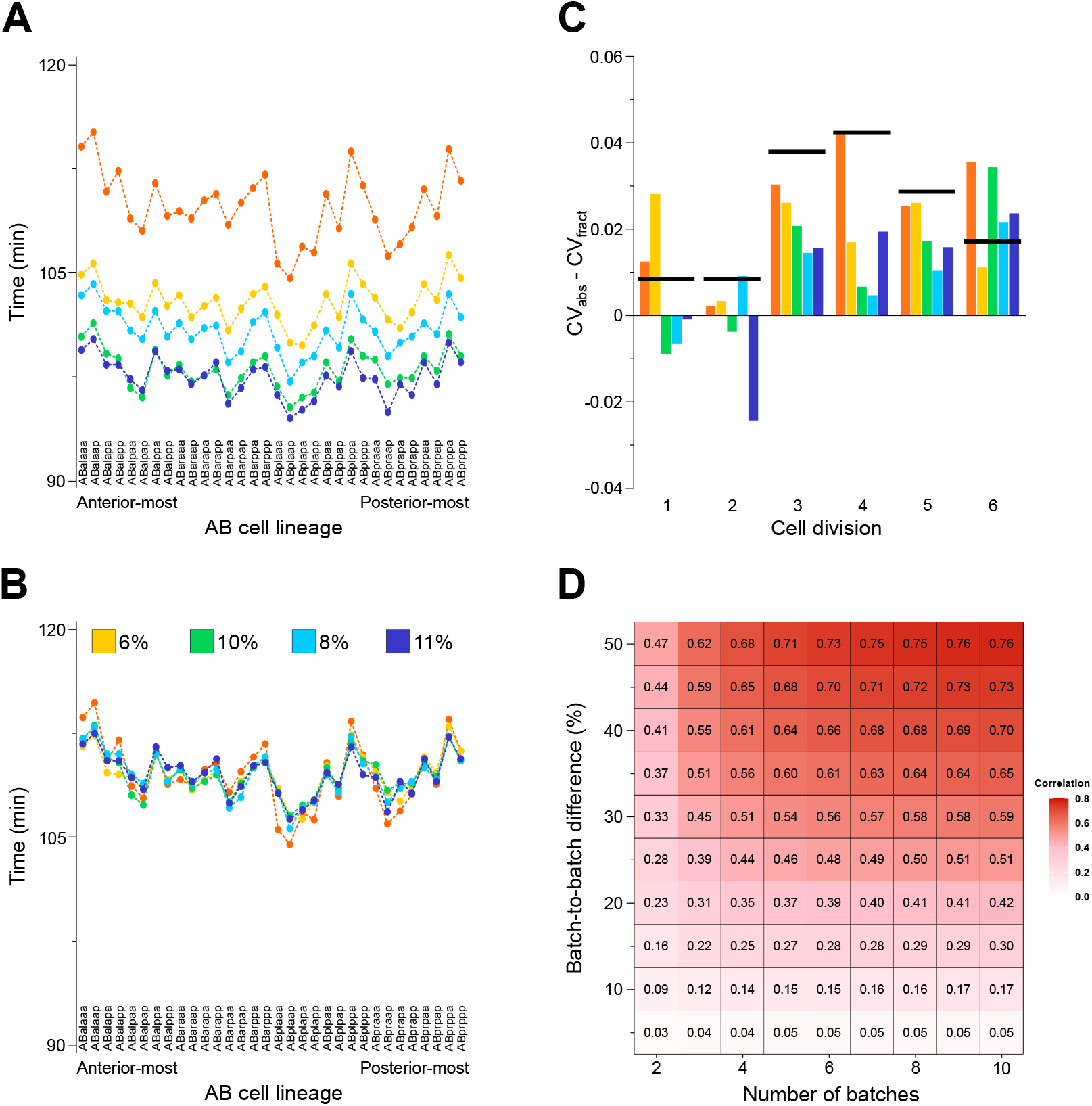
Batch effects can result in false correlations. **(A)** Average birth times of the indicated embryonic cells in the AB lineage of *C. elegans* in five different strains. Each dot is an average birth time of a given cell among the ten individuals that constitute a batch. Each batch corresponds to a strain, as indicated by five different colors. **(B)** The average birth times of the indicated cells in the four faster-developing strains (from panel A) became almost perfectly coincident with the average birth times seen in the slowest strain following linear re-scaling by the indicated coefficients. **(C)** Absolute variability (CV_abs_) of timing of all six analyzed embryonic divisions was greater than fractional variability (CV_fract_) for two (orange and yellow) batches, but not consistently so for the remaining three batches. Black bars indicate values from joint analysis of all five batches as in Figure 1F. **(D)** Correlations within randomly generated timeseries as a function of the number of batches and batching effects. Each square shows the average of 10,000 simulated timeseries.

To examine the potential consequences of batch effects on false-discovery of scaling, we modeled a hypothetical development-like process and a typical way in which the data on temporal unfolding of such processes are collected in real experiments. Briefly, we envisioned a four-stage process, akin to *C. elegans* larval development. To construct the total developmental time of one simulated individual, durations of each of the four constituting stages were set at 10 ± a random value between 0 and 1. If durations of development for all simulated individuals were “collected” in this way, no temporal scaling would be expected because durations of stages are uncorrelated. To recapitulate the fact that data are often collected in batches and then collectively analyzed, developmental timeseries for simulated individuals constructed as above were assigned into batches of the same size with all stage durations in a given batch being multiplied by a “batch effect” constant. As expected, greater number of batches and greater “batch effects” (i.e., batch-to-batch differences) increased false correlations between simulated developmental times (Figure 3D). Interestingly, sample size, tested in the range of 1 to 500, had only modest impact on batch effects (Figure S4C).

The results of our simulation suggest that the inference of scaling may be impacted by batching artifacts. Correlations of ∼0.3-0.4, comparable to those seen in real data (e.g. [20]), can readily arise from batch-to-batch differences of ∼15-20% across ∼4-5 batches (Figure 3D). These conditions are quite typical in real experiments (Figure 3B). Critically, however, batching uniformly inflates all correlations. Thus, if in a given analysis correlations between duration of some sequential processes are higher than others, it may still be possible to conclude that some of these processes scale, while others do not. Then, following the logic described in the previous section, it should be possible to infer shifts in the underlying biological processes, despite the impact of batch effect.

### Correlations within timeseries can be beneficial

Several notable exceptions (Figure 2) notwithstanding, scaling is common in biological timeseries. Whereas we demonstrated that scaling inevitably arises from correlations inherent in continuous biological processes, we also wondered about the consequences of such correlations for the organisms. As a concrete example, we revisited larval development of *C. elegans* that consists of four stages L1-L4. As we demonstrated above, temporal scaling requires that the associations between durations of larval stages in individuals are not random – longer-than-average L1s are more likely to be followed by longer-than-average L2s, etc. Consequently, in a population of developing *C. elegans* larvae, the distribution of durations of total developmental times is going to be overdispersed relative to a distribution in which durations of larval stages L1-L4 are sampled at random from the same empirical set of stage durations (Figure 4A). Practically speaking, more real animals would have both longer and shorter total durations of larval development than in a hypothetical population with the identical mean durations of each larval stage (L1-L4) but without stage-to-stage correlations. A population of animals with overdispersed development times would have an advantage over a population with uncorrelated stage durations if having particularly slow or fast developers is beneficial. Such situations are likely common in nature.

**Figure 4.**
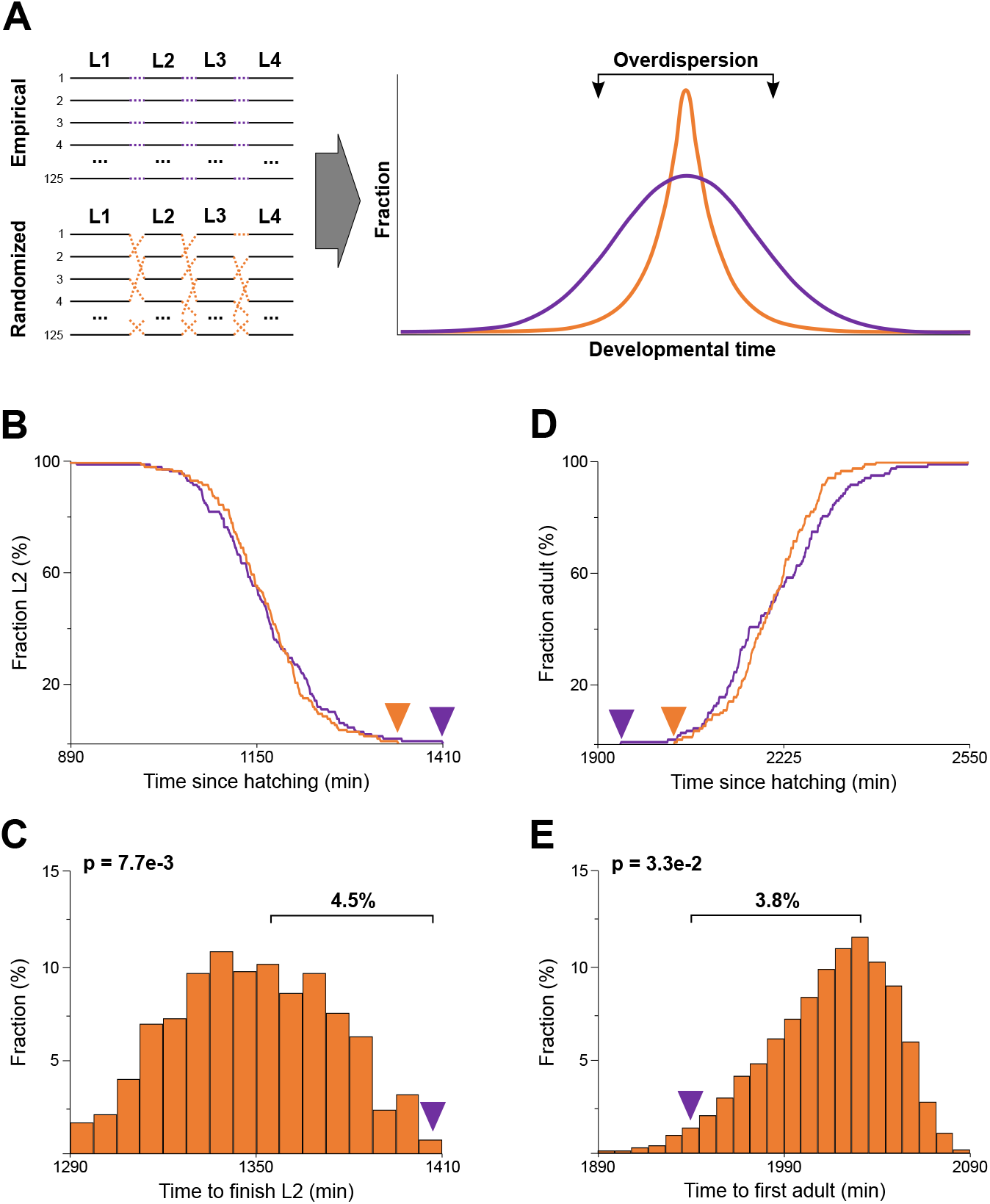
Correlated processes distribute risk more broadly over time. **(A)** The panel on the top left depicts an empirical timeseries consisting of 125 individuals (L1-L4 are four discrete periods). Below is a simulated set of 125 generated by randomly selecting an L1, an L2, an L3, and an L4 from the respective empirical distribution. The empirical (i.e., correlated) timeseries is overdispersed relative to the randomized comparison (right). **(B)** Fraction of individuals remaining in the L2 larval stage as a function of time in the empirical (purple) and one representative randomized population (orange). Arrowheads point to the minute when the last individual in each population exited the L2. **(C)** Distribution (out of 10,000 simulated runs) of times when at least one “individual” in the randomized populations was schmauer-capable (i.e., still in the L2). In the empirical population, the last individual exited the L2 (purple arrowhead) ∼4.5% later than the simulated average. The difference between the empirical and randomized populations is significant (p = 7.7e-3). **(D)** Fraction of individuals achieving adulthood as a function of time in the empirical (purple) and one representative randomized population (orange). Arrowheads point to the minute when the first individual in each population entered adulthood. **(E)** Distribution (out of 10,000 simulated runs) of times when the first “individual” in the randomized populations became an adult. In the empirical population, the first individual became adult (purple arrowhead) ∼3.8% earlier than the simulated average. The difference between the empirical and randomized populations is significant (p = 3.3e-2).

We first examined the benefits of the excess of slow developers. Because the environment is unpredictably variable and habitats are often suboptimal, animals have evolved suitable survival strategies. For example, under unfavorable environmental conditions, *C. elegans* larvae can exit reproductive development to become dauer larvae, resilient morphs that can persist for long periods of time [33, 34]. However, the developmental decision to become dauer can only be made during a relatively narrow time window during the L1 larval stage; if conditions worsen later, becoming dauer is no longer possible [35].

Consider a hypothetical developmental decision, schmauer, identical to dauer except occurring during the L2, before the end of that larval stage (there must be at least two larval stages to test for the consequences of random vs. non-random association between their durations). In a population of 125 worms (data from [16], as analyzed in [20]), the latest individual to finish the L2 did so at ∼1405 minutes post hatching (Figure 4B). In a simulated set (N=125) in which durations of L1 and L2 stages were drawn and assorted at random from the empirical developmental timeseries, the last “simulated worm” to reach the end of the L2 did so ∼62 minutes earlier. At that time, the empirical population had several worms in the L2, therefore still capable of undergoing the schmauer decision. A sudden onset of unfavorable conditions (that could only be survived in schmauer) would be better tolerated by the empirical, overdispersed population because having even one individual still in the L2 stage would allow it to be reestablished, given that *C. elegans* are self-fertilizing hermaphrodites.

To estimate the benefit provided by the overdispersion of developmental time, we computationally generated 10,000 sets of 125 pairs of L1 and L2 stage durations randomly drawn from the empirical set. The last worm to finish the L2 in the empirical population did so ∼4.5% later than the last individual in an average simulated population (Figure 4C). Consequently, the empirical population consisting of 125 worms would have survived the sudden onset of unfavorable conditions better than 99% (9,923 out of 10,000) of simulated populations of the same size. Curiously, all of the remaining 77 simulated populations had all “individuals” finish the L2 by exactly 1405 minutes.

We next considered the benefit of excess of fast developers. Many species live in ephemeral environments which have limited resources that are only sufficient to support a population for a short time. Life in such competitive environments follows the boom-and-bust cycles that consist of arrival, rapid expansion, and collapse that can be followed by migration. In such an environment, the earlier an arriving individual initiates reproduction, the more likely it is to gain advantage over others [36]. Because the natural habitats of *C. elegans* are thought to be ephemeral [37], we compared the distribution of ages at larval-to-adult transition (a proxy for the onset of reproduction) in the empirical population of 125 worms and 10,000 simulated populations of the same size (generated as above, by random sampling and assortment, but this time of four larval stages – L1 through L4). As expected, the fastest developing worm in the empirical population became adult ∼161 minutes earlier than the fastest simulated worm (Figure 4D). Among the simulated sets, only ∼3% (327/10,000) yielded one or more animal that reached adulthood faster than the fastest worm from the empirical population (Figure 4E).

To summarize, internal correlations that are ubiquitous in biological timeseries result in overdispersed distributions of developmental time and, therefore, cause the excess of individuals on the faster and slower extremes. Existence of such individuals effectively broadens the population-wide duration for all biological processes and in this way increases the probability that at least some individuals would be in a given state. In essence, internal correlations in biological processes distribute risk more broadly over time.

## Discussion

Our study offers empirical evidence that durations of discrete stages commonly scale within biological timeseries. Because correlation is the necessary and sufficient condition for this kind of scaling, the same rationale would apply to any quantitative variable that changes over time. We therefore posit that the kind of scaling we described will be observed for a broad range of traits in all organisms.

The scaling we described can be readily diagnosed by comparing variance of absolute and relative durations of stages, an equivalent approach being applicable to any other trait as well. Therefore, scaling can be detected using a simple and transparent statistical test of variance for any quantitative, time-variable data. We propose that the fundamental reason for the correlations that result in scaling is the continuity of biological processes from one stage to another or from one value of a quantitative trait to another. For this reason, failing to observe scaling is the “surprising” result likely indicating a substantial change in the underlying processes. Thereafter more nuanced statistical models (e.g [18, 19]) and empirical studies should be used to identify the causal mechanistic changes.

A downside of a diagnostic test that relies on the inference of correlation is that batching effects could result in false correlations and create the appearance of scaling where none exists. In the case we considered, batching refers to multiple biological replicates of the same experiment being pulled together to increase sample size. Whereas batching of this kind is a concern for accurate inference of both means and correlations, the latter are particularly affected. As our simulations show, modest batch-to-batch variability results in artificial values of correlation that are well in range of those commonly observed in real data. Moreover, increasing the number of batches exacerbates the problem of falsely inferred correlations. A plausible source of batch effects is the variability in experimental conditions that is extrinsic to the intraindividual correlations that are being studied. Potential sources of batch effects could involve differences in physical parameters (temperature, humidity, etc.) or biological states (age, nutrition, etc.). Separating intrinsic from extrinsic variability could be challenging [38], while completely eradicating or even controlling for experimental noise may be practically impossible. For these reasons, the observed extent of scaling is likely to be an overestimate, particularly if the data are collected in batches. While this fact is a potential limitation of our approach, we nevertheless believe that the simple comparison of absolute and fractional variance has considerable utility. As we argue, because scaling is the expected observation, the truly interesting findings are the cases that deviate from it. Batch effects artificially inflate estimates of all correlations equally, making the observation of non-scaling conservative with respect to false-discovery.

Finally, we demonstrated that correlated timeseries have a property that is likely to be beneficial in natural environments. A population of individuals undergoing an internally correlated process will be overdispersed relative to the uncorrelated control. The increased representation of slow and fast individuals stretches the time during which some members of the population are in a particular state, effectively increasing robustness with respect to the environmental fluctuations that are a universal feature of natural habitats.

## Acknowledgements

We are deeply grateful to the colleagues who generously shared advice and the primary data that were analyzed in this study – Z. Bao, D. Charabidze, L. Guignard, W. Keil, P. Keller, P. Lemaire, O. Loudet, K. McDole, J. I. Murray, Y. Sasakura, S. Stern, T. Triphan, S. Uppaluri, F. Vasseur, D. Weigel. The primary data for several studies analyzed here were deposited with the original publications. The data on progeny production in *C. elegans* were collected in the IR laboratory by Erin Aprison. This study would not have been possible without the help of all of the above individuals. We could only hope that all researchers showed the same level of transparency, scholarly engagement, willingness to share. We thank Rick Morimoto for generous hospitality. This work was funded in part by an NSF grant (IOS-1755244) to IR.

## Materials and Methods

### Variability analysis in individualized datasets

The data analyzed in this study were generously shared by multiple colleagues. The data on *C. elegans* locomotory behavior (Figure 1C) were reported by [16] and further analyzed by [20]. The dynamics of *A. thaliana* leaf vegetative growth (Figures 1D, S1A) were reported by [21]. *C. elegans* brood accumulation data (Figure 1E, Table S1) were collected in the IR laboratory; these data were described in [39], but individual-resolved data were not previously released. Timing of cell divisions in early *C. elegans* embryogenesis (Figures 1F, S1B) was obtained from [22-24]. Timing of cell divisions in *P. mamillata* embryogenesis (Figures S1C, S1D) was obtained from [25]. *C. elegans* growth during larval development (Figure 2A) was reported in [26]. The data on duration of development and lifespan in *C. elegans* (Figure 2B) were obtained from [11]. Durations of larval, pupal, and adult stages in *D. melanogaster* (Figure 2C) were reported by [13]. Development time in *C. vicina* and *L. sericata* (Figures S2B, S2C) were obtained from [28]. In each case, individualized timeseries (e.g. distance travelled in 30 min, increase in rosette area per day, time between cell divisions) were extracted from the primary data and the corresponding fractions of total quantities (e.g. ratio of gain in rosette area in a given day to total gain) were calculated. Coefficients of variation of absolute (CV_abs_) and fractional (CV_fract_) sets were calculated and compared.

### Estimation of batching in *C. elegans* embryonic lineage data

The entire dataset containing timing of *C. elegans* embryonic divisions (Figure 1F) was separated into five “batches” corresponding to the five different strains used in the experiments (Table S1). Average time to complete the first four rounds of divisions was calculated for all ABa and ABp descendants in each batch. Distributions of these developmental times were found to be significantly different (Kolmogorov-Smirnov tests) in all pairwise comparisons except Batch 3 vs. Batch 5 (Figure S4A). For all five batches, the average time to complete four divisions was computed (that is, the average of the 32 time-to-completion values per batch). The ratio of Batch 1 average to Batch X average is the batch-normalization coefficient (where X = 2, 3, 4, 5). Batches 2 through 5 were linearly scaled by multiplying individual values of time-to-completion of four divisions (32 values per batch) by the appropriate batch-normalization coefficient (Figure 3B). The data of the six-division lineage (ABalaaaal) presented in Figure 1F were separated into the same five batches each corresponding to a different strain from which the data were collected. Batch properties appeared to be highly similar for the first six division and for the first four divisions (Figure S4B). Absolute (CV_abs_) and fractional (CV_fract_) variabilities in the lineage in which six divisions were considered (ABalaaaal) were then calculated and compared separately for each batch.

### Simulating batch effects on correlations in randomly generated data

We used computational simulation to evaluate the extent to which correlations between stages within timeseries could be affected by batching. We generated simulated data in the following way – each individual’s “development” consisted of four “stages” lasting 10 ± a random value between 0 and 1 of arbitrary time units. Several “individuals” (*N*_*batch_size*_) simulated in this way were assigned to a “batch”; a number of “batches” (*K*_*batches*_) were generated. To simulate batch effects, all values in each “batch” were multiplied by a “batch coefficient” = 1 + *Δ* (*Δ* was selected at random between 0 and a value of *Δ*_*max*_ the range of which is described below). Each combination of three parameter values (*K*_*batches*_, *N*_*batch_size*_, *Δ*_*max*_) yielded correlations between all possible pairs of the four “stages”. An average of these six correlations can serve as a metric of the impact of batch effects. For each combination of the three parameters, we ran 10,000 simulations. The average correlations of these runs are shown in Figure 3D. We tested the following parameter values: *K*_*batches*_ = 2 to 10; *N*_*batch_size*_ = 1, 2, 5, 10, 20, 35, 50, 75, 100, 200 and 500; *Δ*_*max*_ = 0.05 to 0.50 in 0.05 increments. The code used to generate these simulated data is deposited at https://github.com/denisfaer/Faerberg_et_al_2023/blob/main/batching_heatplot.pas

### Randomizing *C. elegans* larval stage duration data

To provide the baseline for comparisons of overdispersion (Figure 4), we generated randomized timeseries of *C. elegans* larval development. An L1, an L2, an L3, and an L4 were randomly selected from the distributions of durations of respective stages in the empirical dataset [20]; four of these stages together produced an “individual”. Each “set” consisted of 125 “individuals”, same as the size of the empirical set. The total of 10,000 randomized sets was generated. For the experiments in Figures 4B, C, we identified the time when the last individual in all sets (one empirical and 10,000 randomized) exited the L2. For the experiments in Figures 4D, E we determined the time when the first individual in all sets transitioned from the L4 to adulthood. p-values were calculated as the fraction of 10,000 randomized sets in which at least one “individual” a) remained in the L2 at the last time when the last animal in the empirical set was still in the L2 (Figure 4C) or b) was already an adult before adults appeared in the empirical set (Figure 4E). The advantage of overdispersion was calculated as percent difference between empirical value and the mean of the corresponding 10,000 randomized sets. The code used to generate these data is deposited at https://github.com/denisfaer/Faerberg_et_al_2023/blob/main/overdisp.pas

