## Supplemental Figures for "Scaling, correlations, and artifacts in biological timeseries"

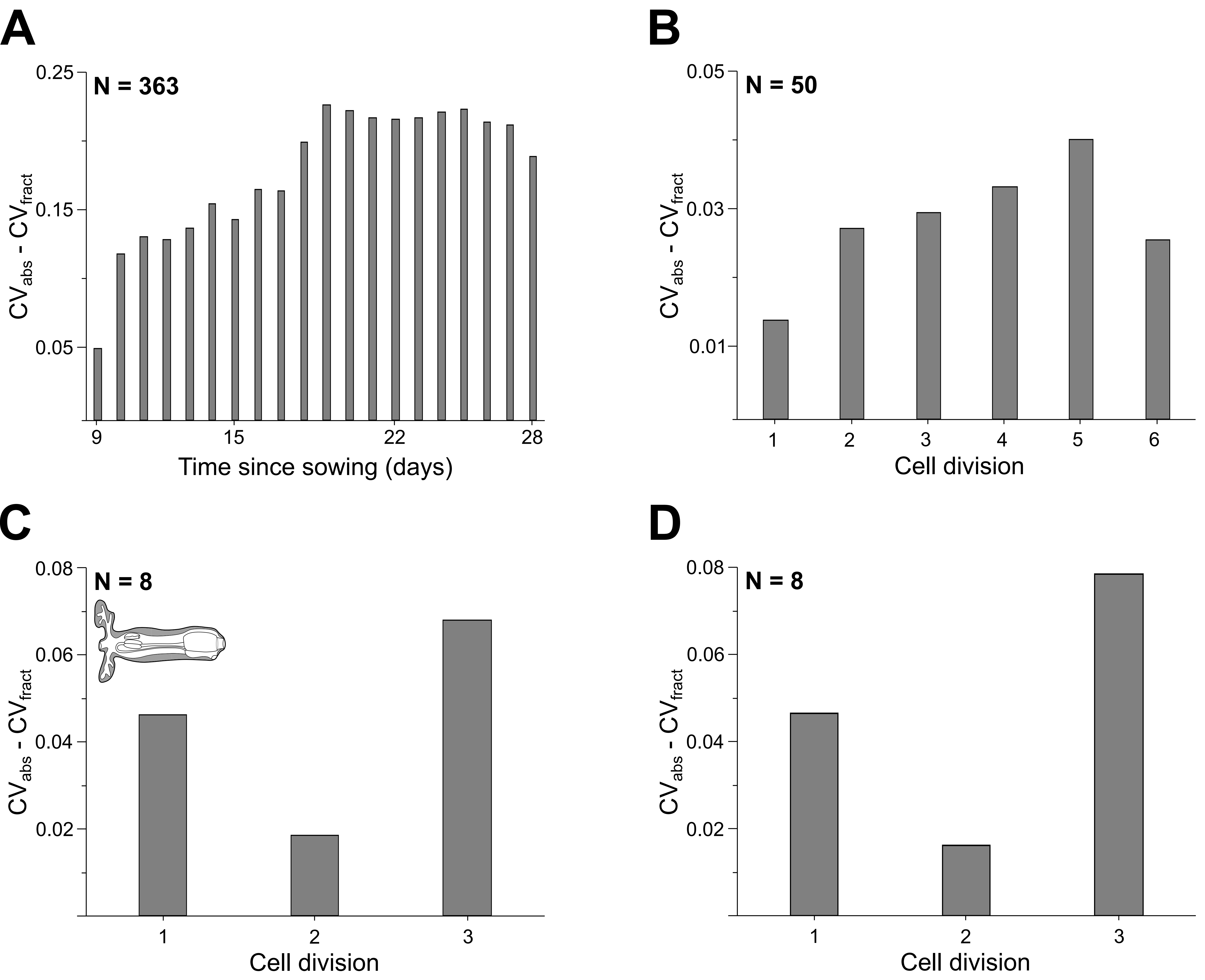
A
N = 363
0.25
0.15
0.05
$CV_{abs}$ - $CV_{fract}$
9
15
22
28
Time since sowing (days)
B
N = 50
0.05
0.03
0.01
$CV_{abs}$ - $CV_{fract}$
1
2
3
4
5
6
Cell division
C
N = 8
0.08
0.06
0.04
0.02
$CV_{abs}$ - $CV_{fract}$
1
2
3
Cell division
D
N = 8
0.08
0.06
0.04
0.02
$CV_{abs}$ - $CV_{fract}$
1
2
3
Cell division

**Figure S1. Additional comparisons of absolute and fractional variability in biological timeseries. Related to Figure 1.**
(A) Absolute ($CV_{abs}$) vs. fractional ($CV_{fract}$) variability in leaf vegetative growth in *A. thaliana* (related to Figure 1D). The data used for this calculation were derived from an independent experimental replicate reported in the same study as the data analyzed in Figure 1D. (B) Absolute ($CV_{abs}$) vs. fractional ($CV_{fract}$) variability in division timing in the ABprppppp embryonic lineage in *C. elegans* (related to Figure 1F). (C) Absolute vs. fractional variability in division timing in the b7.0014*-b8.0027*-b9.0053* lineage in *P. mammillata*. (D) Absolute ($CV_{abs}$) vs. fractional ($CV_{fract}$) variability in the lineage b7.0014_-b8.0027_-b9.0053_ in *P. mammillata*. Sample sizes for each variability analysis are shown in the top left corner of the corresponding bar plot.

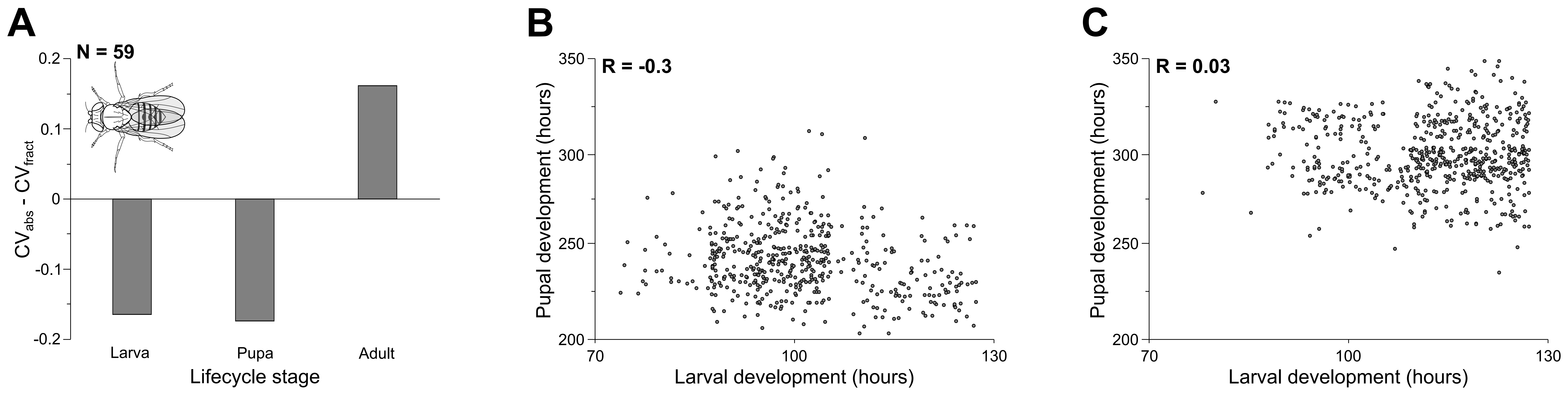
A
N = 59
0.2
0.1
0
-0.1
-0.2
CV_abs - CV_fract
Larva
Pupa
Adult
Lifecycle stage
B
R = -0.3
350
300
250
200
Pupal development (hours)
70
100
130
Larval development (hours)
C
R = 0.03
350
300
250
200
Pupal development (hours)
70
100
130
Larval development (hours)

**Figure S2. Additional evidence that increased fractional variability results from low correlations. Related to Figure 2.**
(A) Absolute ($CV_{abs}$) vs. fractional ($CV_{fract}$) variability of time spent as larva, pupa or adult in *D. melanogaster* females (N=59). (B) Correlation between durations of larval and pupal stages in *L. sericata* (N=502). (C) Correlation between durations of larval and pupal stages in *C. vicina* (N=577).

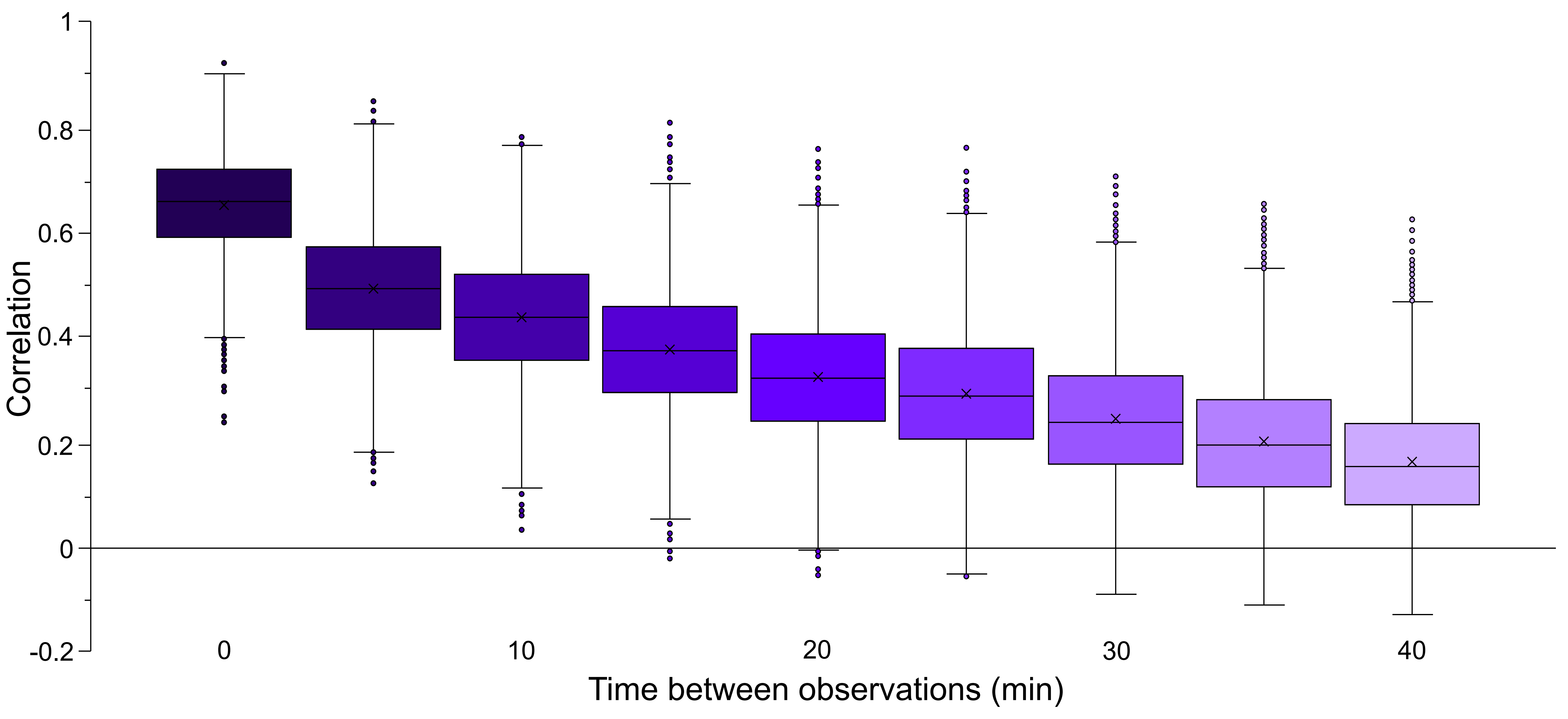

Correlation
1
0.8
0.6
0.4
0.2
0
-0.2
0
10
20
30
40
Time between observations (min)

**Figure S3. Correlations of activity levels decay over time.**
We calculated correlations between locomotory activity of *C. elegans* larvae during consecutive time periods. We selected a random 5-minute period in the L3 larval stage and nine consecutive 5-minute periods. For each of these ten periods, the total distance traveled was computed. This procedure was repeated for each of the 125 worms in the sample. For each comparison of activity during the initial 5-minute period to the nine subsequent ones, correlations were computed using the entire set of worms (N=125). We repeated this procedure 10,000 times to obtain accurate estimates of average correlations. Code used to generate this data is deposited at https://github.com/denisfaer/Faerberg_et_al_2023/blob/main/activity_cumul.pas

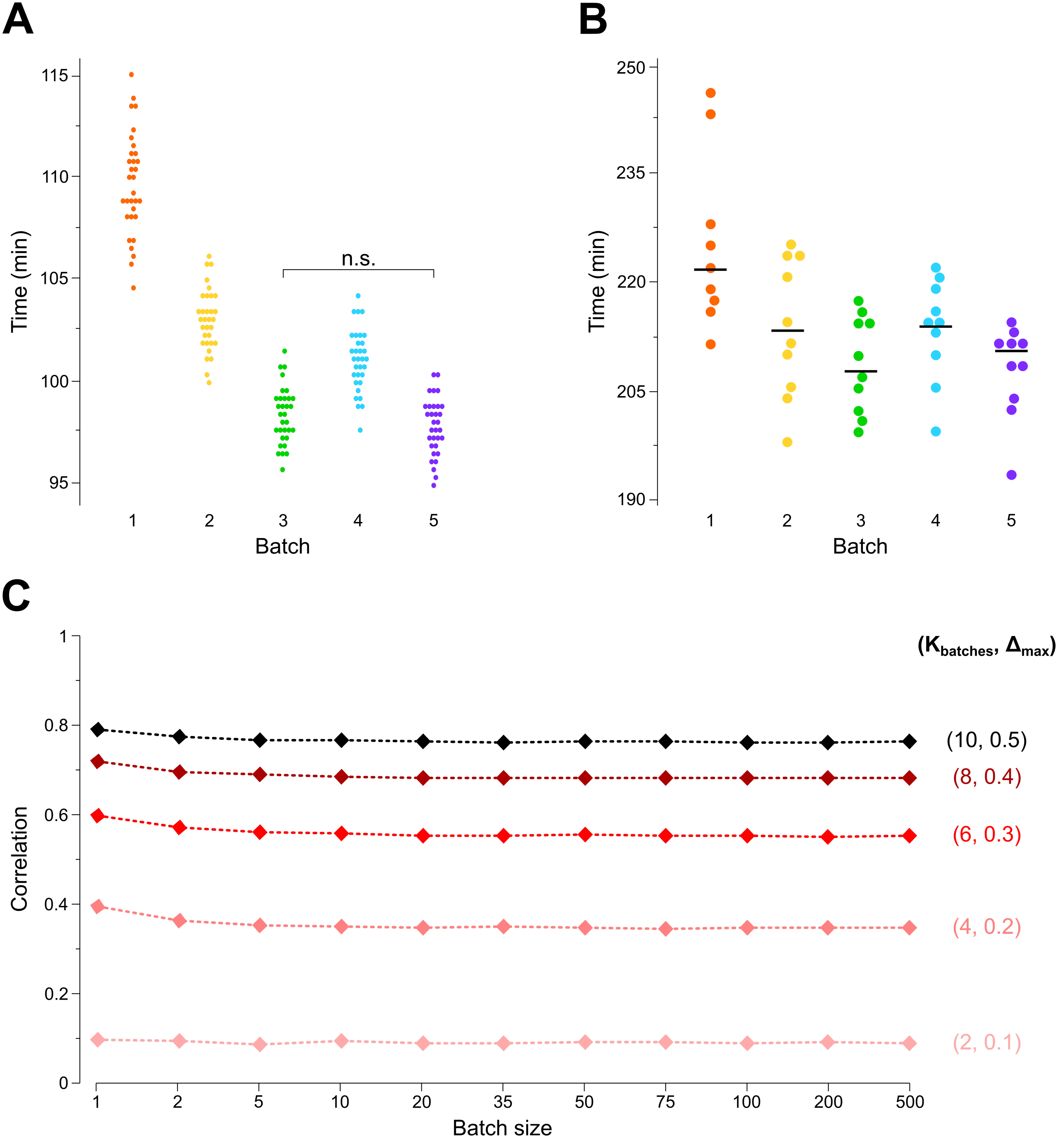

A
Time (min)
115
110
105
100
95
n.s.
1
2
3
4
5
Batch
B
Time (min)
250
235
220
205
190
1
2
3
4
5
Batch
C
Correlation
1
0.8
0.6
0.4
0.2
0
($K_{batches}$, $\Delta_{max}$)
(10, 0.5)
(8, 0.4)
(6, 0.3)
(4, 0.2)
(2, 0.1)
1
2
5
10
20
35
50
75
100
200
500
Batch size

**Figure S4. Batching effects in real and simulated data sets. Related to Figure 3.**
(A) Replotting of the data shown in Figure 3A. Each dot represents the average birth time of the cells resulting from the fourth embryonic division in the ABa and ABp lineages. All pairwise batch comparisons except for the indicate one are significant (Kolmogorov-Smirnov test). (B) Absolute time to the birth of a specific cell (ABalaapaa) in embryos of five different strains (batches). Each dot represents one embryo. Black bars indicate means. (C) The impact of batch size on the artificial correlations introduced by batching. Values of $K_{batches}$ and $\Delta_{max}$ used to construct each curve are shown on the right.
